# PDEparams: Parameter fitting toolbox for partial differential equations in Python

**DOI:** 10.1101/631226

**Authors:** César Parra-Rojas, Esteban A. Hernandez-Vargas

## Abstract

**Motivation:** Partial differential equations (PDEs) is a well-established and powerful tool to simulate multi-cellular biological systems. However, available free tools for validation against data are not established. The `PDEparams` module provides flexible functionality in Python for parameter estimation in PDE models.

**Results:** The `PDEparams` module provides a flexible interface and readily accommodates different parameter analysis tools in PDE models such as computation of likelihood profiles, and parametric bootstrapping, along with direct visualisation of the results. To our knowledge, it is the first open, freely available tool for parameter fitting of PDE models.

**Availability and implementation:** The `PDEparams` module is distributed under the MIT license. The source code, usage instructions and step-by-step examples are freely available on GitHub at github.com/systemsmedicine/PDE_params.

## 1 Introduction

PDE models appear in a wide variety of biological contexts (Anderson *et al.* (2000); Jaeger *et al.* (2004); Reis *et al.* (2016); Hross and Hasenauer (2016)) and, while most available computational tools focus on the numerical integration of PDE models to varying degrees of efficiency and complexity—see, *e.g.*, Guyer *et al.* (2009) or Alnæs *et al.* (2015)—we have not come across general-use implementations incorporating functionality for parameter optimisation with respect to data, and the analysis of parameter identifiability and variability. Moreover, a wide range of models in biology consist of simple equations, in simple spatial domains, and the data available for validation tends to be sparse. To the best of our knowledge, there is no publicly available, open and free-to-use tools for kinetic parameter estimation of PDE models, but only codes for specific examples mainly for ordinary differential equations (ODEs)—*e.g.*, Nguyen and Hernandez-Vargas (2018). Here we present `PDEparams`, a free Python module for parameter fitting in PDE models, and the analysis of parameter estimates in a straightforward, intuitive manner.

## 2 Materials and methods

The `PDEparams` module is not only meant to work when all variables in the system are observed, but also in the more realistic case when data are available for only one or a few of them; additionally, we accommodate the case when the observed quantity corresponds to a function of the variables, rather than their raw values.

### 2.1 The PDEmodel object

The main component of the module is the `PDEmodel` object. As the input, the user provides the data (as a pandas (McKinney *et al.* (2010)) `DataFrame`), along with the PDE model (as a function of the state vector), the initial condition (as a function of the coordinate vector), and the lower and upper bounds for the unknown parameters—the estimation is carried out using the Differential Evolution (DE) algorithm (Storn and Price (1997)), which performs constrained optimisation. Other arguments include the parameter names (used for tables and plots, defaults to ‘parameter 1’, ‘parameter 2’, …); the number of variables in the system (defaults to 1); the number of spatial dimensions—for completeness, the module can also handle ODEs, for which this value should be set to zero (defaults to 1); the number of replicates, defined as the number of different measurements per space-time coordinate (defaults to 1); the observed variable if not all are observed (all assumed observed by default); and the function to apply to the raw variables (defaults to None, and raw outputs are used).

The constructed object contains a time array and spatial grid—to which the initialising functions have been applied—for integration of the model. These and all other vector operations are carried out using NumPy (Van Der Walt *et al.* (2011)).

### 2.2 Best fit parameters

After construction of the `PDEmodel` object, parameters that provide the best fit between model and data— within their specified bounds—can be obtained by simply running the `fit()` method. If no argument is provided, the function to be minimised will be the mean squared error (MSE). Other options include: i) the root mean squared error; ii-iii) the mean (and root mean) squared logarithmic error; iv) the mean absolute error; and v) the median absolute error. The errors are computed using scikit-learn’s (Pedregosa *et al.* (2011)) built-in functions; the integration of the model itself is handled with SciPy *(Jones et al.* (01)), as is the DE optimisation.

When `fit()` is run, the best parameters and the lowest error are added as attributes to the `PDEmodel` object. The former are also printed to the screen.

### 2.3 Likelihood profiles

Likelihood profiles (Raue *et al.* (2009)) are computed for each parameter by fixing its values on a predefined grid and re-estimating all the rest. This is done running the `likelihood_profiles()` method. When no argument is given, grid of size 100 is used as the default; for different grid sizes, the argument `npoints` may be used. If the best fit parameters have already been obtained, the error to be used for the likelihood profiles will match the one originally used with the `fit()` method. If not, the default mean squared error will be used. During estimation, a progress bar (da Costa-Luis *et al.* (2019)) is displayed on the screen.

As a result, a `DataFrame` with the parameter values and their corresponding error for each of the parameters is added as an attribute to the `PDEmodel` object. These results can then be used internally or be exported as, *e.g.*, a `.csv` file.

### 2.4 Bootstrapping

Parametric bootstrapping is carried out by randomly choosing one replicate per space-time coordinate and re-estimating all parameters in multiple rounds. This is done with the method `bootstrap()`; the number of rounds is controlled by the argument `nruns` which, if not given, is assumed to be 100—note that, if only one measurement per space-time coordinate exists in the data, bootstrapping amounts to simply running the `fit()` method multiple times, and therefore has no effect. If the best fit parameters have already been obtained, the error to be used for bootstrapping will match the one originally used with the fit() method. If not, the default mean squared error will be used. During estimation, a progress bar (da Costa-Luis *et al.* (2019)) is displayed on the screen.

As a result of the procedure, two `DataFrame` objects are added as attributes to the `PDEmodel` object: i) a statistical summary of the parameter values—printed to the screen; and ii) the raw results. These can then be used internally or be exported as, *e.g.*, a `.csv` file.

### 2.5 Visualisations

Within the module, the likelihood profiles and the bootstrapping results can be directly visualised, respectively, using the methods `plot_profiles()` and `plot_bootstrap()`. These use Matplotlib (Hunter (2007)) and Seaborn (Waskom *et al.* (2017)). If the best fit parameters have already been obtained, they will be highlighted in the plots, as shown in Fig. 1.

**Figure 1:**
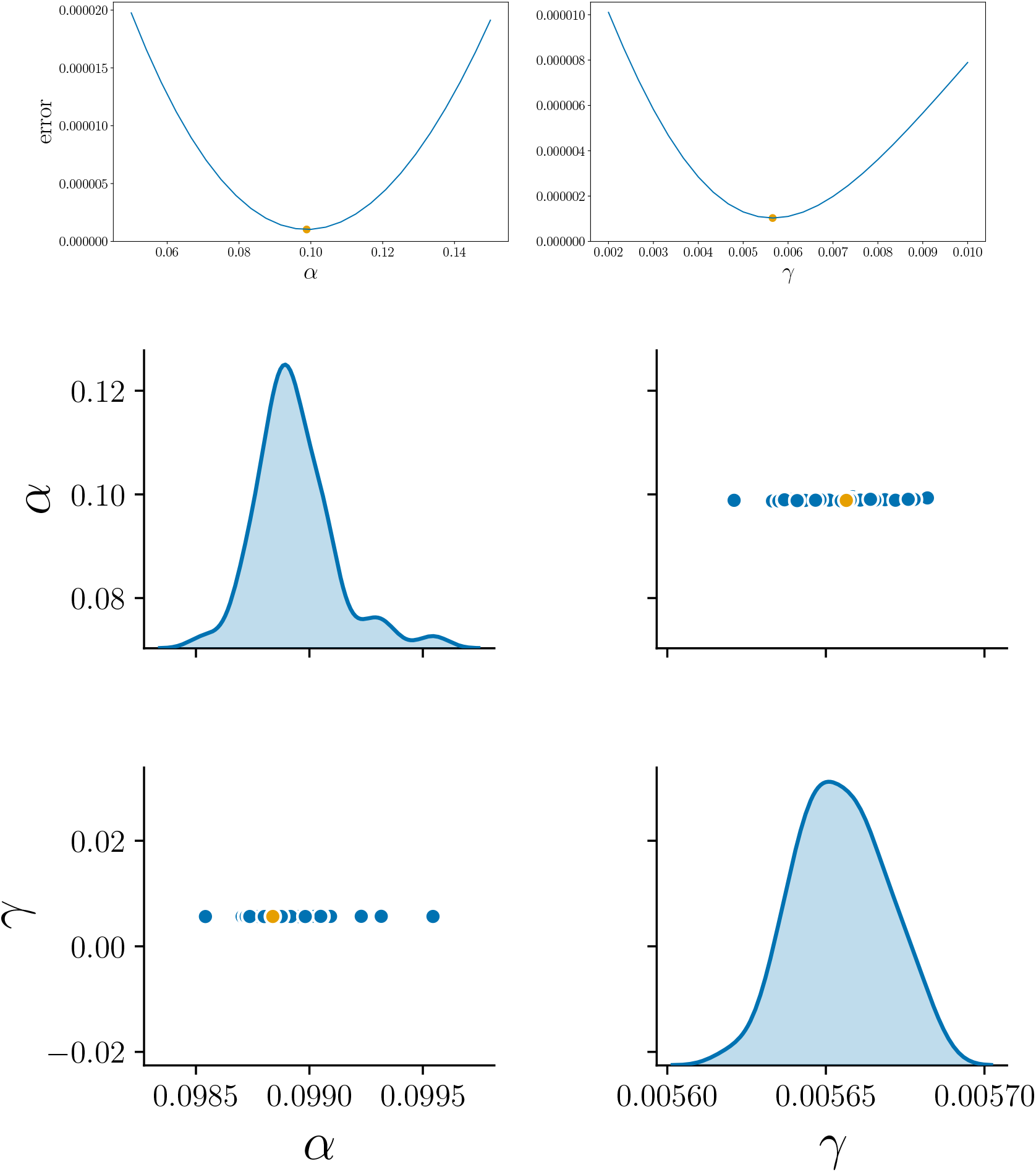
Likelihood profiles (top) and bootstrapping results (bottom) for the estimation of *α* and *γ* from Eq. (1) using only the data for *m*. Visualisations obtained, respectively, with the `plot_profiles()` and `plot_bootstrap()` functions of `PDEparams`. Best fit parameters are shown in orange. Nominal values: *α* = 0.1, *γ* = 0.005

## 3 Example: tumour growth

As an example use case, we take the two-dimensional, three-variable PDE model from *Anderson et al.* (2000), describing the dynamics of the invasion of host tissue by tumour cells:

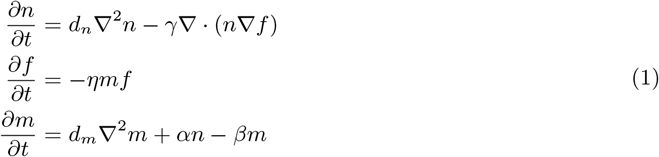

Here, *n* corresponds to the tumour cells, *f* to the host tissue, and *m* to matrix-degradative enzymes associated with the tumour cells.

We start by integrating the model using the same parameters as in the paper—*d*_*n*_ = 0.001, *d*_*m*_ = 0.001, *η* = 10, *α* = 0.1, *γ* = 0.005, and *β* = 0—and the same initial condition for *n* and *m*; the initial condition for the host tissue is heterogeneous and arbitrarily chosen—cf. Fig. 8 from the paper. After this, we generate artificial data by sampling the resulting dynamics on a 25 × 25 spatial grid at times *t* = 1, 2,…, 15. We then use `PDEparams` to estimate the values of *α* and *γ*. The results are summarised in Fig. 1, for the case when only the data for *m* are used for the estimation—*i.e.*, *n* and *f* are assumed unobserved. The full step-by-step example is provided in the Supplementary Material, and is available in the module repository as a Jupyter notebook.

## 4 Conclusions

`PDEparams` is the first free module for the validation of PDE models against data and the analysis of their parameter estimates.

