## Supplemental for "PDEparams: Parameter fitting toolbox for partial differential equations in Python"

**GitHub: github.com/systemsmedicine/PDE_params**

---

### Case study: tumour growth

#### Importing packages

##### Others

In [1]:
```
# Used to read the data file
import pandas as pd

# Used in the definition of the model and its initial condition
import numpy as np

# Used for plots other than those generated by the parameter estimation module
import matplotlib.pyplot as plt
%matplotlib inline
```

##### The module itself

In [2]:
```
import PDEparams as pde
```

#### Defining the model

We use the two-dimensional model of tumour growth from Eq. (5) of Anderson *et al.* (2000)[1]:

$$\frac{\partial n}{\partial t} = d_n \nabla^2 n - \gamma \nabla \cdot (n \nabla f)$$
$$\frac{\partial f}{\partial t} = -\eta m f$$
$$\frac{\partial m}{\partial t} = d_m \nabla^2 m + \alpha n - \beta m$$

Note that in the original simulations, the parameter $\beta$ is set to zero. We do the same here.

##### Boundary conditions

We use no-flux boundary conditions in the 2D domain defined by the data. Eqs. (6)-(7) of Anderson *et al.* (2000) translate to

$$\frac{\partial n}{\partial x} = \frac{\gamma n}{d_n} \frac{\partial f}{\partial x}$$

at the $x$ boundaries; and equivalently for $y$.

Note that the equation for $f$ does not have spatial derivatives, and therefore no boundary conditions. In the code below, the derivatives of $f$ entering the equation for $n$ are computed directly, and then plugged into the condition for the first-order derivative of $n$.

##### Second-order derivatives of $n$ and $m$

In practice, if we consider the variable $u$ ($n$ or $m$) as a discrete 1D vector $(u_0, u_1, \ldots)$, then we can obtain the central differences

$$h\, u_i' = u_{i+1} - u_{i-1}$$
$$h^2\, u_i'' = u_{i+1} + u_{i-1} - 2u_i$$

where $h = dx$, $u' = \partial u/\partial x$, and $u'' = \partial^2 u/\partial x^2$. If we have a boundary condition $u_0' = \sigma$, and we take an imaginary point $u_{-1}$ to the left of our domain so that

$$h^2\, u_0'' = u_1 + u_{-1} - 2u_0$$

at the boundary, we obtain from the first-order difference $u_{-1} = u_1 - h\sigma$. Therefore, in our case

$$h^2 n_0'' = 2n_1 - h\gamma n_0 f_0'/d_n - 2n_0$$
$$h^2 m_0'' = 2m_1 - 2m_0$$

The same idea applies in the 2D case to the four boundaries.

---

[1] Anderson, A.R., Chaplain, M.A., Newman, E.L., Steele, R.J. and Thompson, A.M., "Mathematical modelling of tumour invasion and metastasis", *Computational and Mathematical Methods in Medicine* ***2(2)***, pp.129-154 (2000).

In [3]:

```
def TumourGrowth(z, t, grid, alpha, gamma):

    '''The input z corresponds to the current state of the system, and it's a flattened vector in both the num
ber
    of outputs and the spatial dimensions.

    t is the current time.

    grid is the spatial grid of the model.

    aN and aL correspond to the unknown parameters.
    '''

    # Known parameter values (same as in the paper)

    dN = 1e-3
    dM = 1e-3
    eta = 10.

    '''Here we obtain both functions by reshaping the input: we divide it into 3 portions (number of outputs)
    of whatever shape the spatial grid has (this is what the -1 stands for)'''

    N, F, M = z.reshape(3,-1)

    '''Now we reshape both functions using the actual dimensions of the spatial grid.

    The shape of the spatial grid array is given by:

                        (number of points in dim 1, number of points in dim 2, ..., ndims)

    with ndims the number of spatial dimensions. This is due to the fact that we have a grid of

                        (number of points in dim 1)x(number of points in dim 2)

    elements, but each element is ndims-dimensional. Therefore, we must take the first "ndims" elements of the
    shape of the grid, ignoring the last one, to reconstruct the shape of each function. The slice with [:-1]
    stands for "all elements up to and excluding the last one".
    '''

    N = N.reshape(grid.shape[:-1])

    F = F.reshape(grid.shape[:-1])

    M = M.reshape(grid.shape[:-1])

    '''The grid has the form

                (x0,y0), (x0,y1), ...
                (x1,y0), (x1,y1), ...

    So that:
        x0 = first element of the first element of the first row of the grid: grid[0,0,0]
        x1 = first element of the first element of the second row of the grid: grid[1,0,0]
        y0 = second element of the first element of the first row of the grid: grid[0,0,1]
        y1 = second element of the second element of the first row of the grid: grid[0,1,1]
    '''

    dx = grid[1,0,0]-grid[0,0,0]
    dy = grid[0,1,1]-grid[0,0,1]

    # We initialise the spatial derivatives we need as empty arrays of the same shape of our N and L functions

    dNdx = np.empty_like(N)
    dNdy = np.empty_like(N)
    dNdxx = np.empty_like(N)
    dNdyy = np.empty_like(N)

    dMdxx = np.empty_like(M)
    dMdyy = np.empty_like(M)

    #First-order derivatives

    # np.gradient(array, axis) returns centred 1st-order differences between values along x (rows, axis=0)
    # or y (columns, axis=1), plus forward/backward differences for the left/right boundaries
    dFdx = np.gradient(F, axis=0)/dx
    dFdy = np.gradient(F, axis=1)/dy


    # Left boundary. In the grid array, first row, all columns: [0,:]
    dNdx[0,:] = gamma*N[0,:]*dFdx[0,:]/dN

    # Away from the x-boundary. In the grid array, second to penultimate row, all columns: [1:-1,:]
    dNdx[1:-1,:] = np.gradient(N, axis=0)[1:-1,:]/dx

    # Right boundary. In the grid array, last row, all columns: [-1,:]
    dNdx[-1,:] = gamma*N[-1,:]*dFdx[-1,:]/dN

    # Bottom boundary. In the grid array, all rows, first column: [:,0]
    dNdy[:,0] = gamma*N[:,0]*dFdy[:,0]/dN

    # Away from the y-boundary. In the grid array, all rows, second to penultimate columns: [:,1:-1]
    dNdy[:,1:-1] = np.gradient(N, axis=1)[:,1:-1]/dy
```

```
    # Right boundary. In the grid array, all rows, last column: [:,-1]
    dNdy[:,-1] = gamma*N[:,-1]*dFdy[:,-1]/dN

    # Second-order derivatives

    dFdxx = np.gradient(dFdx, axis=0)/dx
    dFdyy = np.gradient(dFdy, axis=1)/dy

    # Left boundary. In the grid array, first row, all columns: [0,:]
    dNdxx[0,:] = (2.0*N[1,:] - dx*gamma*N[0,:]*dFdx[0,:]/dN - 2.0*N[0,:])/dx**2

    # Away from the x-boundary. In the grid array, second to penultimate row, all columns: [1:-1,:]
    # np.diff(array, order, axis) is used to compute 2nd-order differences between values along x, or rows (ax
is=0)
    dNdxx[1:-1,:] = np.diff(N,2,axis=0)/dx**2

    # Right boundary. In the grid array, last row, all columns: [-1,:]
    dNdxx[-1,:] = (2.0*N[-2,:] - dx*gamma*N[-1,:]*dFdx[-1,:]/dN - 2.0*N[-1,:])/dx**2

    # Bottom boundary. In the grid array, all rows, first column: [:,0]
    dNdyy[:,0] = (2.0*N[:,1] - dy*gamma*N[:,0]*dFdy[:,0]/dN - 2.0*N[:,0])/dy**2

    # Away from the y-boundary. In the grid array, all rows, second to penultimate columns: [:,1:-1]
    # np.diff(array, order, axis) is used to compute 2nd-order differences between values along y, or columns
 (axis=1)
    dNdyy[:,1:-1] = np.diff(N,2,axis=1)/dy**2

    # Right boundary. In the grid array, all rows, last column: [:,-1]
    dNdyy[:,-1] = (2.0*N[:,-2] - dy*gamma*N[:,-1]*dFdy[:,-1]/dN - 2.0*N[:,-1])/dy**2

    # Now the same for M

    dMdxx[0,:] = (2.0*M[1,:] - 2.0*M[0,:])/dx**2
    dMdxx[1:-1,:] = np.diff(M,2,axis=0)/dx**2
    dMdxx[-1,:] = (2.0*M[-2,:] - 2.0*M[-1,:])/dx**2

    dMdyy[:,0] = (2.0*M[:,1] - 2.0*M[:,0])/dy**2
    dMdyy[:,1:-1] = np.diff(M,2,axis=1)/dy**2
    dMdyy[:,-1] = (2.0*M[:,-2] - 2.0*M[:,-1])/dy**2


    # Adding the rest of the terms to obtain the time derivatives

    dNdt = dN*(dNdxx + dNdyy) - gamma*(dNdx*dFdx + dNdy*dFdy + N*(dFdxx + dFdyy))
    dFdt = -eta*M*F
    dMdt = dM*(dMdxx + dMdyy) + alpha*N

    # We put the three time derivatives side by side

    dzdt = np.array([dNdt, dFdt, dMdt])

    return dzdt.reshape(-1) # We return a completely flattened version of the total time derivative
```

Here we specify the functions that define the initial condition for each variable—Eq. (9) of Anderson *et al.* (2000).

For this example, we used an arbitrarily chosen heterogeneous cellular matrix to generate the data, as in Fig. 8 of the paper.

In [4]:
```
def initial_N(z, centre=[0.5,0.5], rad=0.1):
    epsilon = 2.5e-3
    x, y = z

    r = np.sqrt((x - centre[0])**2 + (y - centre[1])**2)

    if r <= rad:
        return np.exp(-r**2/epsilon)

    return 0.

# Homogeneous ECM (only for reference)
def initial_F(z):
    return 1. - 0.5*initial_N(z)

# Heterogeneous ECM
def initial_F2(z):
    return (1. - initial_N(z, rad=0.5) - 0.5*initial_N(z, centre=[0.8,0.8], rad=0.2)
            - 0.8*initial_N(z, centre=[0.2,0.2], rad=0.1) - 0.6*initial_N(z, centre=[0.05,0.6], rad=0.3)
            - 0.3*initial_N(z, centre=[0.6,0.1], rad=0.05) - initial_N(z, centre=[0.6,0.6], rad=0.5)
            - initial_N(z, centre=[0.8,0.4], rad=0.5) - initial_N(z, centre=[0.9,0.2], rad=0.5)
            - initial_N(z, centre=[0.3,0.8], rad=0.7))

def initial_M(z):
    return 0.5*initial_N(z)
```

### Using `PDEparams` to estimate parameters

First, we load the data from the `.csv` file.

The data consist of 3 replicates, and have been generated using parameter values $\alpha = 0.1$, $\gamma = 0.005$, as in the paper.

The columns are, in order: $t$, $x$, $y$, $n$, $f$, $m$.

In [5]:
```
df = pd.read_csv('TumourGrowthData.csv')

df.head()
```

Out[5]:

| | 0 | 1 | 2 | 3 | 4 | 5 |
|---|---|---|---|---|---|---|
| **0** | 1.0 | 0.0 | 0.020408 | 0.001154 | 1.001187 | 0.000670 |
| **1** | 1.0 | 0.0 | 0.020408 | 0.000828 | 0.998376 | 0.000000 |
| **2** | 1.0 | 0.0 | 0.020408 | 0.000000 | 0.998738 | 0.001314 |
| **3** | 1.0 | 0.0 | 0.061224 | 0.001059 | 1.000115 | 0.000000 |
| **4** | 1.0 | 0.0 | 0.061224 | 0.000886 | 0.999843 | 0.000075 |

##### Constructing the `PDEmodel` object.

The inputs are

**Required:**

1. The data table `df`.
2. The model `TumourGrowth`.
3. The list of initial condition functions.
4. The bounds for the parameter values.

**Optional:**

1. The parameter names.
2. The number of variables: 3. **(Default is 1, this needs to be provided in this case)**
3. The number of spatial dimensions: 2. **(Default is 1, this needs to be provided in this case)**
4. The number of replicates in the data: 3. **(Default is 1, this needs to be provided in this case)**
5. The indices of the measured variables. In this case, the default `None`, since we have data for all 3 variables.
6. The function to apply to the output. In this case, the default `None`, since our data is directly $n$, $f$ and $m$.

In [6]:
```
my_model = pde.PDEmodel(df, TumourGrowth, [initial_N, initial_F2, initial_M],
                        bounds=[(0.05, 0.15), (0.002,0.01)], param_names=[r'$\alpha$', r'$\gamma$'],
                        nvars=3, ndims=2, nreplicates=3, obsidx=None, outfunc=None)
```

Let us plot the initial condition.

In [7]:
```
X = my_model.space[:,:,0] # all x values
Y = my_model.space[:,:,1] # all y values

plt.contourf(X,Y,my_model.initial_condition[0], cmap='magma')
plt.colorbar()
plt.title(r'$n(x,y,0)$')
plt.xlabel(r'$x$')
plt.ylabel(r'$y$')
plt.show()

plt.contourf(X,Y,my_model.initial_condition[1], cmap='magma')
plt.colorbar()
plt.title(r'$f(x,y,0)$')
plt.xlabel(r'$x$')
plt.ylabel(r'$y$')
plt.show()

plt.contourf(X,Y,my_model.initial_condition[2], cmap='magma')
plt.colorbar()
plt.title(r'$m(x,y,0)$')
plt.xlabel(r'$x$')
plt.ylabel(r'$y$')
plt.show()
```

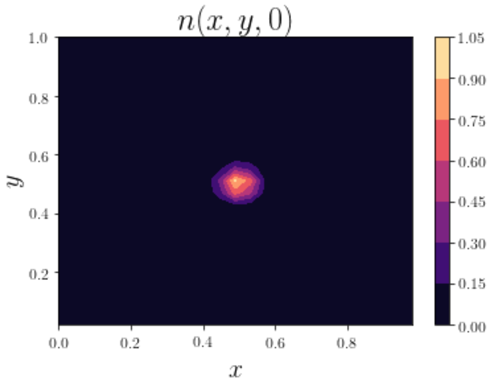


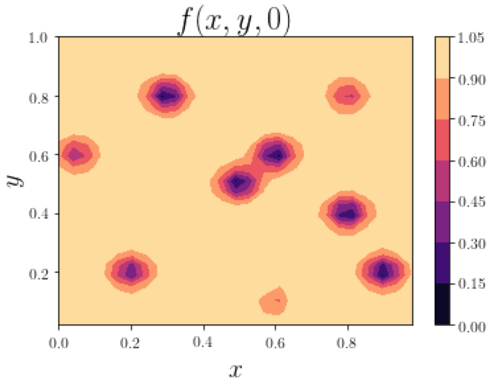


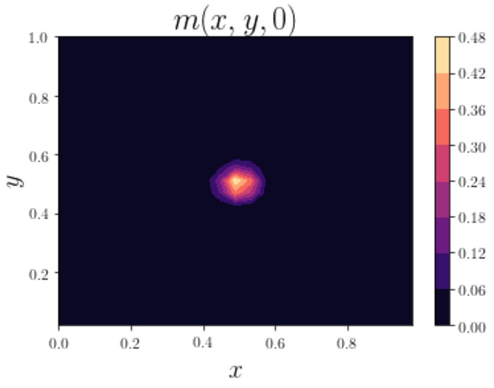


**Finding the best fit parameters.**

In [8]:
```
%%time
my_model.fit()
```

```
   $\alpha$  $\gamma$
0  0.103305  0.005293
CPU times: user 1min 42s, sys: 1.98 s, total: 1min 44s
Wall time: 1min 39s
```

In [9]:
```
my_model.best_params
```

Out[9]:

| | $\alpha$ | $\gamma$ |
|---|---|---|
| **0** | 0.103305 | 0.005293 |

In [10]:
```
my_model.best_error
```

Out[10]:
```
1.3730150561063116e-05
```

##### Likelihood profiles

We use a grid of 25 points per parameter.

**Note:** if you see a "widget not found" message, just ignore it; a progress bar will appear when you run the cell below.

In [13]:
```
%%time
my_model.likelihood_profiles(npoints=25)
```

```
CPU times: user 1h 21min 4s, sys: 8min 19s, total: 1h 29min 23s
Wall time: 1h 2min 39s
```

Visualisation

In [15]:
```
my_model.plot_profiles()
```

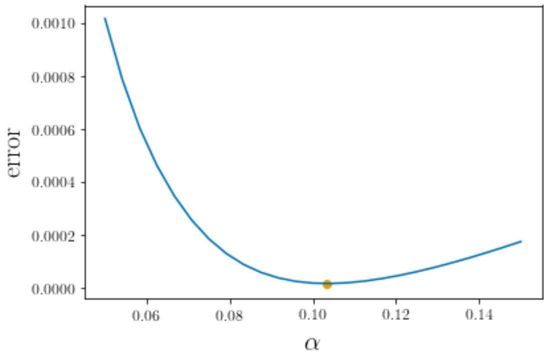


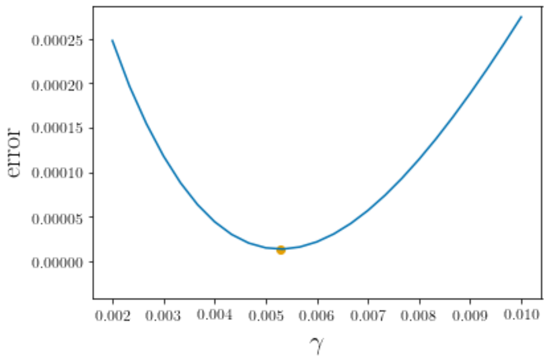


##### Bootstrapping

We use 50 repetitions.

**Note:** if you see a "widget not found" message, just ignore it; a progress bar will appear when you run the cell below.

In [16]:
```
%%time
my_model.bootstrap(nruns=50)
```

```
        $\alpha$   $\gamma$
count  50.000000  50.000000
mean    0.103383   0.005287
std     0.000092   0.000007
min     0.103137   0.005275
25%     0.103337   0.005283
50%     0.103400   0.005286
75%     0.103443   0.005291
max     0.103549   0.005304
CPU times: user 1h 13min 4s, sys: 2min 24s, total: 1h 15min 28s
Wall time: 1h 8min 14s
```

The summary

In [17]:
```
my_model.bootstrap_summary
```

Out[17]:

| | $\alpha$ | $\gamma$ |
|---|---|---|
| **count** | 50.000000 | 50.000000 |
| **mean** | 0.103383 | 0.005287 |
| **std** | 0.000092 | 0.000007 |
| **min** | 0.103137 | 0.005275 |
| **25%** | 0.103337 | 0.005283 |
| **50%** | 0.103400 | 0.005286 |
| **75%** | 0.103443 | 0.005291 |
| **max** | 0.103549 | 0.005304 |

Visualisation

In [19]:
```
my_model.plot_bootstrap()
```

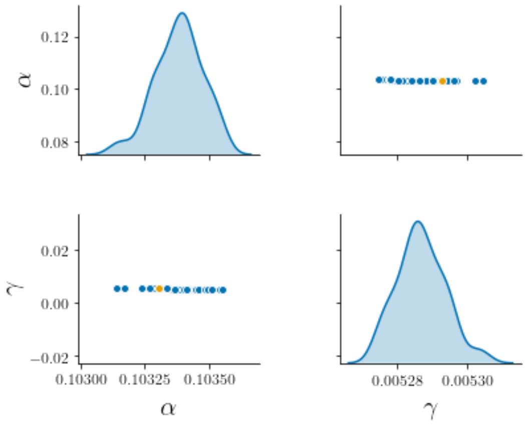


### Only one observed variable

Let us suppose now that we only have data for $m$.

In [20]:
```
df2 = df[df.columns[[0,1,2,5]]] # n and f, which we are ignoring, are columns 3 and 4 in the data table

df2.head()
```

Out[20]:

| | 0 | 1 | 2 | 5 |
|---|---|---|---|---|
| **0** | 1.0 | 0.0 | 0.020408 | 0.000670 |
| **1** | 1.0 | 0.0 | 0.020408 | 0.000000 |
| **2** | 1.0 | 0.0 | 0.020408 | 0.001314 |
| **3** | 1.0 | 0.0 | 0.061224 | 0.000000 |
| **4** | 1.0 | 0.0 | 0.061224 | 0.000075 |

##### Constructing the `PDEmodel` object.

The inputs are the same as before, except for the data—now `df2` instead of `df` —and `obsidx` , which specifies that we only have data for the 3rd variable in the system. Starting from zero, the corresponding index is 2. Since we only have one observed variable, `obsidx` can be either a number or a list of one element.

In [21]:
```
my_model2 = pde.PDEmodel(df2, TumourGrowth, [initial_N, initial_F2, initial_M],
                         bounds=[(0.05, 0.15), (0.002,0.01)], param_names=[r'$\alpha$', r'$\gamma$'],
                         nvars=3, ndims=2, nreplicates=3, obsidx=[2], outfunc=None)
```

##### Finding the best fit parameters.

In [22]:
```
%%time
my_model2.fit()
```

```
   $\alpha$  $\gamma$
0  0.098852  0.005657
CPU times: user 1min 14s, sys: 2.89 s, total: 1min 17s
Wall time: 1min 8s
```

```
In [23]: my_model2.best_params
```

Out[23]:

| | $\alpha$ | $\gamma$ |
|---|---|---|
| **0** | 0.098852 | 0.005657 |

```
In [24]: my_model2.best_error
```

Out[24]: 1.0316723898609223e-06

#### Likelihood profiles

**Note:** if you see a "widget not found" message, just ignore it; a progress bar will appear when you run the cell below.

```
In [25]: %%time
my_model2.likelihood_profiles(npoints=25)
```

```
CPU times: user 49min 40s, sys: 3min 25s, total: 53min 5s
Wall time: 42min 29s
```

Visualisation

```
In [27]: my_model2.plot_profiles()
```

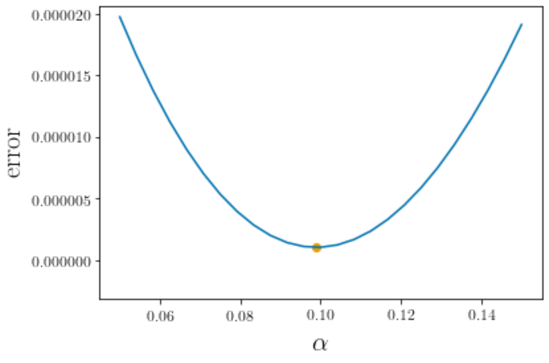


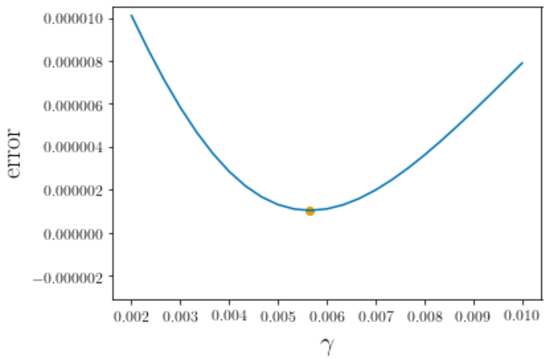


#### Bootstrapping

**Note:** if you see a "widget not found" message, just ignore it; a progress bar will appear when you run the cell below.

```
In [28]: %%time
my_model2.bootstrap(nruns=50)
```

```
        $\alpha$   $\gamma$
count  50.000000  50.000000
mean    0.098954   0.005656
std     0.000135   0.000013
min     0.098674   0.005623
25%     0.098862   0.005648
50%     0.098947   0.005656
75%     0.099022   0.005664
max     0.099387   0.005684
CPU times: user 58min 36s, sys: 1min 58s, total: 1h 34s
Wall time: 54min 28s
```

The summary

In [29]:
```
my_model2.bootstrap_summary
```
Out[29]:

| | $\alpha$ | $\gamma$ |
|---|---|---|
| **count** | 50.000000 | 50.000000 |
| **mean** | 0.098954 | 0.005656 |
| **std** | 0.000135 | 0.000013 |
| **min** | 0.098674 | 0.005623 |
| **25%** | 0.098862 | 0.005648 |
| **50%** | 0.098947 | 0.005656 |
| **75%** | 0.099022 | 0.005664 |
| **max** | 0.099387 | 0.005684 |

Visualisation

In [31]:
```
my_model2.plot_bootstrap()
```
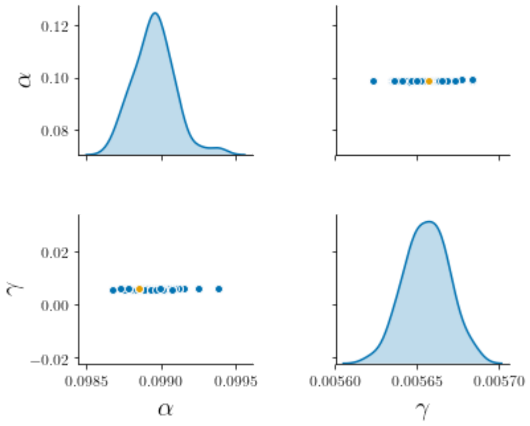


### Only a function of the variables is observed

Finally, we consider the case when it's not the raw outputs of the system that are observed, but rather a function of them. In this case, let us assume that the observed quantity is

$$\mathcal{F}(n, f, m) = n^2 - m$$

In [32]:
```
df3 = df[df.columns[[0,1,2]]] # we take only the space-time coordinates

# We add the new function
df3[r'$\mathcal{F}$'] = df['3']*df['3'] - df['5'] # n and m are, respectively, columns 3 and 5 in the data tab
le

df3.head()
```
Out[32]:

| | 0 | 1 | 2 | $\mathcal{F}$ |
|---|---|---|---|---|
| **0** | 1.0 | 0.0 | 0.020408 | -6.687299e-04 |
| **1** | 1.0 | 0.0 | 0.020408 | 6.859981e-07 |
| **2** | 1.0 | 0.0 | 0.020408 | -1.314136e-03 |
| **3** | 1.0 | 0.0 | 0.061224 | 1.122511e-06 |
| **4** | 1.0 | 0.0 | 0.061224 | -7.427586e-05 |

##### Constructing the `PDEmodel` object.

The inputs are the same as before, except for the data—now `df3` —and `outfunc` , which specifies the function to be applied to the outputs before computing the error. Note that, even if the output function uses only 2 of the variables, we need to pass the full output vector as an argument (see below). In this case, `obsidx` is ignored, but we set it to `None` anyway.

In [33]:
```
def F(z):
    n, f, m = z

    return n**2-m
```
In [34]:
```
my_model3 = pde.PDEmodel(df3, TumourGrowth, [initial_N, initial_F2, initial_M],
                         bounds=[(0.05, 0.15), (0.002,0.01)], param_names=[r'$\alpha$', r'$\gamma$'],
                         nvars=3, ndims=2, nreplicates=3, obsidx=None, outfunc=F)
```

**Finding the best fit parameters.**

In [37]:
```
%%time
my_model3.fit()
```

```
   $\alpha$  $\gamma$
0  0.099206  0.005681
CPU times: user 1min 46s, sys: 1.7 s, total: 1min 47s
Wall time: 1min 44s
```

In [38]:
```
my_model3.best_params
```

Out[38]:

| | $\alpha$ | $\gamma$ |
|---|---|---|
| **0** | 0.099206 | 0.005681 |

In [39]:
```
my_model3.best_error
```

Out[39]:
```
1.106811988396949e-06
```

**Likelihood profiles**

**Note:** if you see a "widget not found" message, just ignore it; a progress bar will appear when you run the cell below.

In [40]:
```
%%time
my_model3.likelihood_profiles(npoints=25)
```

```
CPU times: user 1h 5min, sys: 3min 13s, total: 1h 8min 13s
Wall time: 58min 3s
```

Visualisation

In [42]:
```
my_model3.plot_profiles()
```

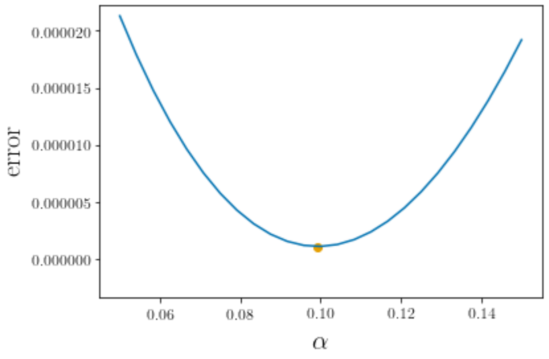


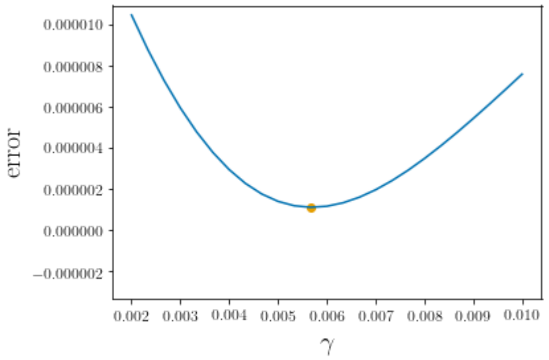


**Bootstrapping**

**Note:** if you see a "widget not found" message, just ignore it; a progress bar will appear when you run the cell below.

In [43]:
```
%%time
my_model3.bootstrap(nruns=50)
```

```
         $\alpha$   $\gamma$
count  50.000000  50.000000
mean    0.099367   0.005695
std     0.000141   0.000015
min     0.099032   0.005660
25%     0.099296   0.005686
50%     0.099364   0.005694
75%     0.099442   0.005702
max     0.099672   0.005729
CPU times: user 1h 6min 39s, sys: 2min 1s, total: 1h 8min 40s
Wall time: 1h 2min 20s
```

The summary

In [44]:
```
my_model3.bootstrap_summary
```

Out[44]:

| | $\alpha$ | $\gamma$ |
|---|---|---|
| **count** | 50.000000 | 50.000000 |
| **mean** | 0.099367 | 0.005695 |
| **std** | 0.000141 | 0.000015 |
| **min** | 0.099032 | 0.005660 |
| **25%** | 0.099296 | 0.005686 |
| **50%** | 0.099364 | 0.005694 |
| **75%** | 0.099442 | 0.005702 |
| **max** | 0.099672 | 0.005729 |

Visualisation

In [46]:
```
my_model3.plot_bootstrap()
```

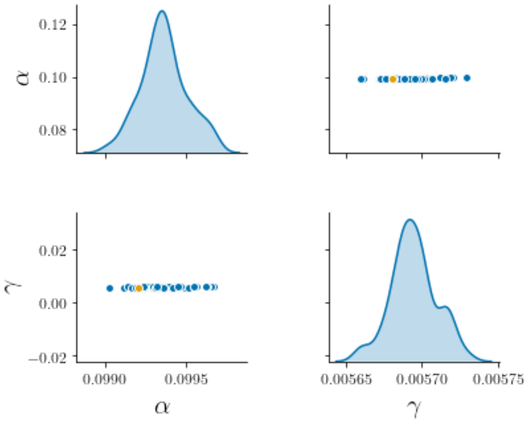
